# Myosin Filaments of Vertebrate Skeletal and Cardiac Muscle are Highly Similar, but not Identical

**DOI:** 10.64898/2026.02.16.706210

**Authors:** Pouria Gholami Tilko, Hosna Rastegarpouyani, Behrouz Ghazi Esfahani, Jose Renato Pinto, Kenneth A. Taylor

## Abstract

Striated muscles consist of two filament types, one composed mostly of actin and the other composed mostly of myosin^1,2^. Actin filaments are highly similar across different muscle types and species both invertebrate and vertebrate^3^. Myosin filaments of vertebrate striated muscle are quite homogeneous in structure having identical lengths, governed by the giant protein titin^4,5^, and rotational symmetries while varying mostly in the isoforms of its proteins. Conversely, myosin filaments from invertebrate striated muscle are highly heterogeneous in multiple ways even within a single organism. Myosin filaments from vertebrate cardiac muscle have been shown to be highly similar in structure between mice and humans^6-8^. Conversely, thick filaments from the highly specialized insect indirect flight muscle have been shown to be highly variable in structure^9-14^. Here we used the drug mavacamten to stabilize a myosin head conformation known as the interacting heads motif in a fast skeletal muscle of rabbits, a highly studied model system. We show that the structure of relaxed rabbit skeletal muscle thick filaments is highly similar to those of relaxed human and mouse cardiac muscle, differing primarily in the positioning of some domains of myosin binding protein C vis-à-vis titin. In the context of the very different structures from indirect flight muscle, the result highlights different solutions to the same problems, control of muscle force and the requirements of endothermy, the internal generation of heat. In mammals, thick filaments are poised for varying levels of myosin activation^15^, while indirect flight muscle is poised for narrowly defined, high frequency contraction. In mammals, endothermy is a continuous problem; in insects, endothermy is primarily necessary for flight^16^.

## Introduction

Vertebrate skeletal muscle constitutes the largest tissue in the animal body, making up 40%–50% of its weight^1^. Striated muscle’s fundamental structural unit is the sarcomere, which is made up of alternating, interdigitated arrays of thin, actin-containing filaments and thick, myosin-containing filaments^2^. The structure of striated muscle actin filaments is well defined varying mostly in the isoform of actin binding proteins while retaining a consistent helical structure^3^. The detailed structure of the striated muscle myosin filaments is only now emerging, boosted by the recent rapid progress in cryoelectron microscopy (cryoEM) image quality from direct electron detectors and image processing software^17^. The first striated muscle thick filaments visualized at subnanometer resolution where protein secondary structure is resolved were isolated from the indirect flight muscles of several insects^9,11,13,14^. Although similar in some details, these thick filaments all had easily visualized differences when reconstructed under relaxing conditions of high ATP and low calcium concentrations. The myosin filaments of vertebrate cardiac muscle have recently been reported showing in detail the arrangement of key proteins like myosin, titin and cardiac myosin binding protein C (cMyBP-C)^6-8^. A structure of thick filaments from vertebrate skeletal muscle at comparable resolution has not been reported.

The precise contraction and relaxation of muscles are crucial for the effective functioning of both cardiac and skeletal muscles, enabling functions like blood circulation, feeding, defense and posture maintenance among many uses. This mechanical force is produced through the interaction between the thick filament motor protein myosin and the thin filament protein actin using chemical energy derived from ATP hydrolysis. Across all vertebrate skeletal and cardiac muscles, the thick filaments consistently measure ∼1.6 μm in length and contain ∼300 myosin dimers^18^.

In addition to myosin, the key proteins that constitute vertebrate thick filaments are titin and MyBP-C. These alone define the three major parts of the vertebrate thick filament relative to the M-line/bare zone in the middle: the proximal zone (P-zone), the central zone (C-zone), and the distal zone (D-zone). The C-zone is distinguished by the presence of MyBP-C^19^. The P-zone and D-zone have other proteins specific to them. Additional constituents such as obscurin^20^ and myomesin^21^ populate the bare zone and are involved in maintaining the ordered, hexagonal arrangement of thick filaments.

Fast myosin binding protein C (fMyBP-C) is a 128 kDa protein predominantly expressed in fast-twitch skeletal muscle. It is characterized by seven immunoglobulin-like and three fibronectin-like domains, an M-domain, and a proline/alanine-rich region. While overall structural similarity has been predicted between fMyBP-C and cardiac myosin binding protein C (cMyBP-C) due to their shared modular domain structure, specific differences exist. cMyBP-C has an extra N-terminal Ig domain (C0), a 28-residue insert in the C5 Ig module and four phosphorylation sites in its M-domain. Unlike cMyBP-C, fMyBP-C has no reported phosphorylation sites. Mutations in fMyBP-C are linked to skeletal muscle diseases like distal arthrogryposis^22^, affecting muscle contractility and force generation.

Previous studies indicate that fMyBP-C plays a role in modulating the speed, force, and power of fast skeletal muscle contraction^23^. High-resolution information on the organization of cMyBP-C in cardiac thick filaments has been published. Whether fMyBP-C shares this detailed structural similarity remains unconfirmed^24^.

## Results

### Structure and Arrangement of Myosin C-zone at 10Å resolution

We conducted a single-particle, 3D reconstruction of the 430Å repeating unit found in the C-zone of native, frozen-hydrated rabbit psoas myosin filaments. This reconstruction, deposited in the Electron Microscopy Data Bank (EMD-74497), achieved a global resolution of ∼10Å (Supplemental Figure 1). We used the drug mavacamten, developed for treating cardiac muscle disease^25^, to stabilize the myosin heads^26^. Mavacamten binds to a pocket between the N-terminal and converter domains of the myosin head^27^ and was used in two recent cardiac thick filament reconstructions^6,7^. We compare our structure primarily with Dutta et al. because both were obtained by single particle methods. Where a contrast with the *in situ* structure is needed, the comparison with Tamborini et al.^6^ and Chen et al.^8^ is provided.

**Figure 1.**
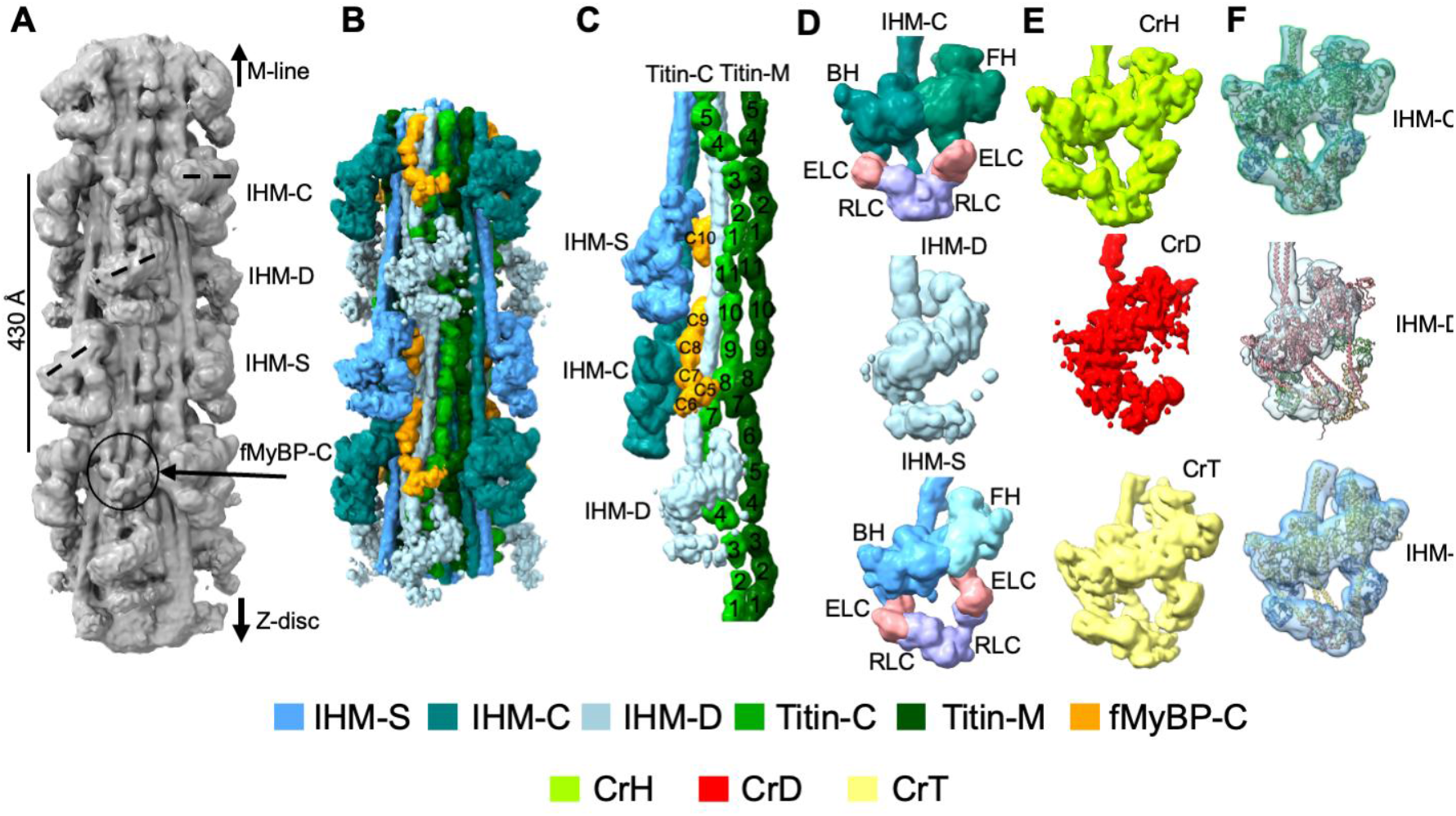
Single-particle 3D reconstruction of the vertebrate skeletal C-zone. Color scheme shown at the bottom is retained in all figures. (A) 3D reconstruction of seven crowns of heads showing three types of myosin head arrangements (IHM-C, IHM-D, and IHM-S), fast skeletal myosin binding protein C (fMyBP-C), and titin. (B) The middle five-crown segment of the seven-crown map from panel A after post-processing using DeepEMhancer, segmented to identify the myosin heads and tails, fMyBP-C molecules (orange), and two titin strands (Titin-C, bright green; Titin-M, dark green). (C) Side view of an asymmetric unit of the reconstruction, oriented to optimize visibility of all components. fMyBP-C domains C5–C10 and titin-M and titin-C super-repeat domains 1–11 are numbered. (D) Structure of IHMs, with their two light chains. ELC, essential light chain (pink), RLC, regulatory light chain (purple). (E) The structures of the three IHM types of the cardiac thick filament, CrH (horizontal), CrD (disordered), and CrT (tilted), are depicted in green, red, and yellow, respectively (compare with Fig. 1D). (F) Atomic models of cardiac IHMs (EMD-15353) were docked into our skeletal IHMs’ 3-D density maps to facilitate a direct comparison of structural conservation. See also Supplementary Video 1.

We first identify the different components in the skeletal thick filament. Then, we will compare these components to those found in cardiac thick filaments, highlighting their similarity and differences.

### Myosin heads

Myosin head pairs were distinctly identified as interacting-head motifs (IHMs), organized into three unique crowns: IHM-D, IHM-C, and IHM-S, using the terminology adopted by Chen et al.^8^ (Fig. 1A-C). These crowns were arranged in triplets, forming an asymmetric unit with a 430Å axial repeat. The filament structure bears no resemblance to a quasi-helical array particularly if the arrangement of myosin tails is also considered. The overall positions of these IHMs along with the thick filament agree with previous studies in cardiac thick filaments^6-8^.

Within each 430Å triplet, the IHM-D density appeared weak and noisy, suggesting significant mobility (Fig. 1D). The IHM-S and IHM-C densities were strong and ordered but differed in orientation: IHM-S was tilted, while IHM-C was nearly horizontal (rotated 20° clockwise from IHM-S). The orientation variation between different IHMs is identical to observations in cardiac muscles^6-8^. Azimuthal rotations within a triplet were 32° from IHM-D to IHM-S and 16° from IHM-C to IHM-S, with a 72° rotation between triplets. All three IHMs of the C-zone repeat showed good fit with the corresponding cardiac thick filament IHMs even to the relative disorder of IHM-D (Fig. 1D-F) and CrD of cardiac muscle^7^.

### Myosin tails

We refer to the arrangement of the myosin α-helical coiled-coil tails as myosin tail layers. Each myosin tail layer consists of myosin tails offset axially by three crowns, 430Å. To deconvolve the complex tail network, we extended the reconstruction to 19 crowns and used color to distinguish the three myosin tail layers segmented from the thick filament backbone (Fig. 2A). The map’s resolution is sufficient to display the individual trajectories of full-length myosin tails that form the filament’s structural core. For all three IHMs, the distal 3/4 of the tail, the light meromyosin segment, pack tightly together with displacements from a straight-line showing complementarity when tails are offset by three crowns (Fig. 2B-E). When just the tails are shown, the proximal 1/4 of the myosin tail, the S2 segment, is clearly separated from the rest of the myosin tail layer but only for IHM-C and IHM-S. The amount of S2 that is free for IHM-D appears about 1/3^rd^ as long (Fig. 2E). The significance of these lengths is that they define the tether that constrains myosin heads when searching for appropriately positioned F-actin subunits for strong myosin binding. However, an interpretation is complex because titin domains appear to be responsible for the separation of myosin tails for IHM-C and IHM-S (see below).

**Figure 2.**
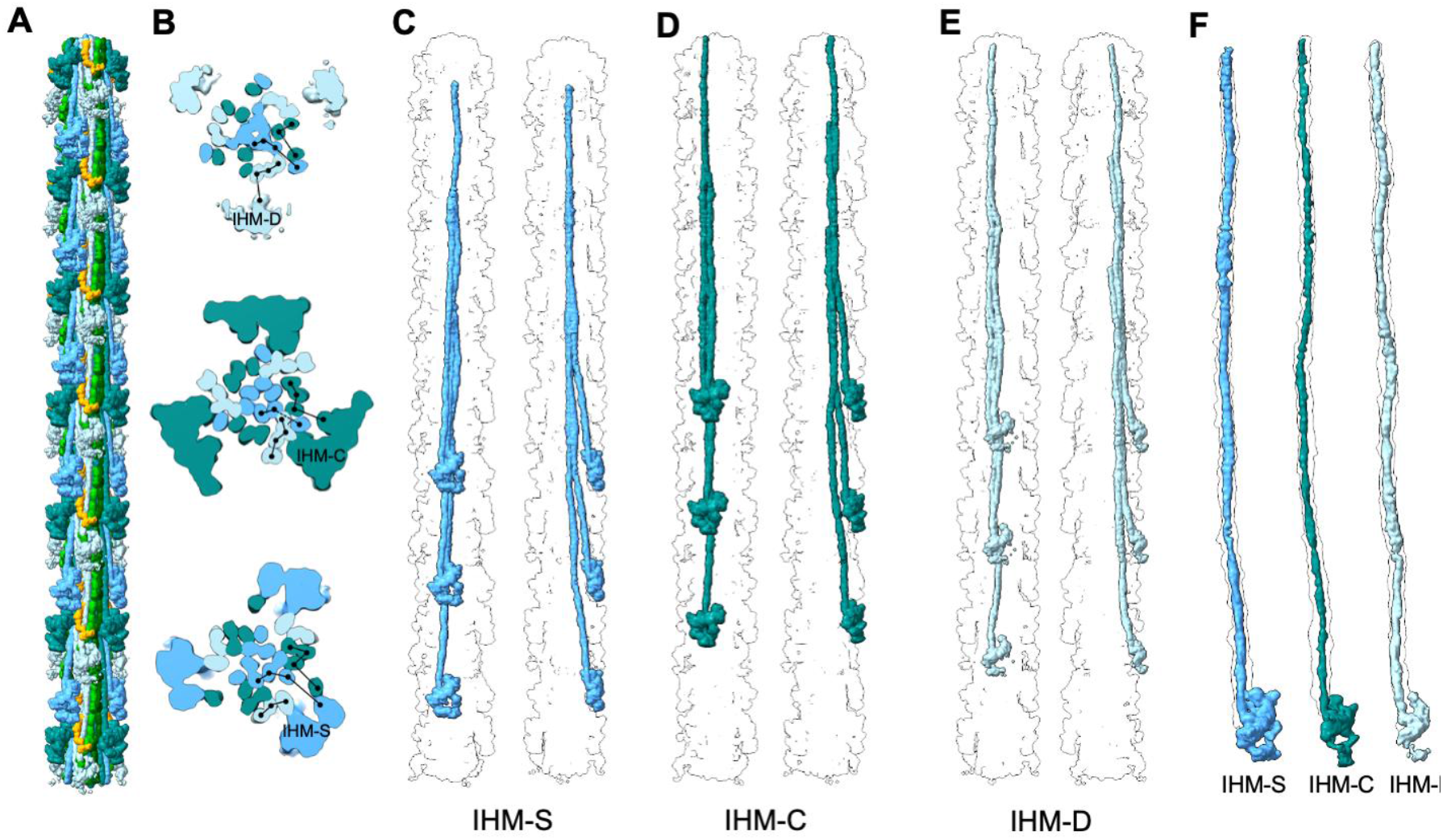
| Myosin tail network in skeletal thick filament C-zone. (A) The generated 19 crown map needed to show complete myosin tail layers illustrating the intricate interaction network within the filament C-zone. (B) Cross-sectional slices of the reconstruction (viewed toward the M-line) showing the organization of the myosin tails at crowns IHM-D, IHM-C, and IHM-S. Black lines indicate the association of individual tails with the separate myosin tail layers. (C-E) Density maps show segmented myosin tails and heads from the three distinct crowns axially offset by 430Å that form the myosin tail layers. The two views have an azimuthal 90º separation. They are designated as (C) IHM-S, (D) IHM-C, and (E) IHM-D. Note that the proximal S2 part of the myosin tail is almost three crowns in length in IHM-S and IHM-C whereas that of IHM-D is only about one crown in length. These patterns are similar to those found in cardiac thick filaments. (F) Fitting the three segmented tails of IHM-C, IHM-S, and IHM-D into the cardiac myosin tails from the human cardiac map model (EMD_40468). See also Supplementary Video 2.

IHM-S tails course deep into the filament’s interior, where they assemble into a tightly packed structure (Fig. 2B,C). They are the only tails that reach the center of the filament, which lacks any space that might accommodate additional proteins. The arrangement of myosin tails in thick filaments of invertebrates produces a hollow core that can accommodate different amounts of the protein paramyosin^9,10^. The space at the center of the vertebrate thick filament is much smaller^7,8^. The IHM-S tail assembly forms the central, load-bearing core of the filament backbone. This architecture is analogous to the tilted crowns (CrT) observed in human cardiac filaments^7^. By being in this central, protected position, the IHM-S myosin tails are possibly greater sensors of tension on the thick filament than other myosin tails, which may be more sensitive to other factors, such as changes in titin or MyBP-C.

IHM-C tails form an intermediate layer between the IHM-S inner core and the IHM-D tails positioned on the outermost surface (Fig. 2B,D). This spatial arrangement mimics the path of tails from the “horizontal crowns” (CrH) in cardiac muscle, which also occupy an intermediate position known to be modulated by MyBP-C interactions^7^. Functionally, this intermediate localization places the IHM-C tails and their associated heads in a position to be regulated by MyBP-C^19^.

The third population, IHM-D, comprises tails that are the most peripheral, forming a loosely packed outer sheath that envelops the other two tail populations (Figs. 2B, E). This path is analogous to that of tails from the “disordered crowns” (CrD) in cardiac muscle^7^. The fact that the myosin heads of IHM-D (CrD) are disordered, structurally poises them to be the first responders to activating signals, such as calcium release activating the thin filament. Being largely disordered (activated) even in the presence of large quantities of mavacamten, suggests that IHM-D heads are unlikely to be recruited by various thick filament activating mechanisms^15^. The outermost position of the IHM-D myosin makes them the most accessible population for protein turnover. Their intimate association with MyBP-C and titin positions them as key players in the integrated regulatory network of the sarcomere^28^.

The distinct paths of these three myosin populations create a radial functional gradient from the filament’s axis to its surface, progressing from deep storage (IHM-S) to a modulated reserve (IHM-C) to a readily available active pool (IHM-D). This trilayered structure provides the physical basis for the “hierarchy of control” and the “three levels of myosin activation” previously described for cardiac muscle^7,8^, enabling the fine-tuned, graded recruitment of motors necessary for efficient muscle function.

To test the hypothesis of conserved architecture, we performed a fitting experiment where the segmented density maps of the three skeletal tail paths (IHM-S, IHM-C, and IHM-D) were docked into the Cryo-EM map of the human cardiac thick filament^7^ (Fig. 2F). The fit demonstrates that each skeletal tail population occupies a position and adopts a conformation that is nearly identical to its counterpart in the cardiac filament.

### Organization of titin in the C-zone

The foundational element of the thick filament backbone in the C-zone is a paired set of titin strands. Two distinct, largely parallel titin strands, titin-C and titin-M, are situated along the filament’s longitudinal axis in each asymmetric unit (Fig. 3A). Titin interactions with the three myosin tail layers in each asymmetric unit is quite variable and mostly confined to myosin’s α-helical coiled-coiled tail domain. The paired titin strands define a highly ordered, super-repeat of the C-zone (Fig. 3B). This length precisely corresponds to the fundamental 430Å periodicity of the myosin heads on the thick filament surface and the regular spacing of associated regulatory proteins, such as MyBP-C, underscoring titin’s role as a molecular ruler or blueprint for thick filament assembly^4,29-31^.

**Fig 3.**
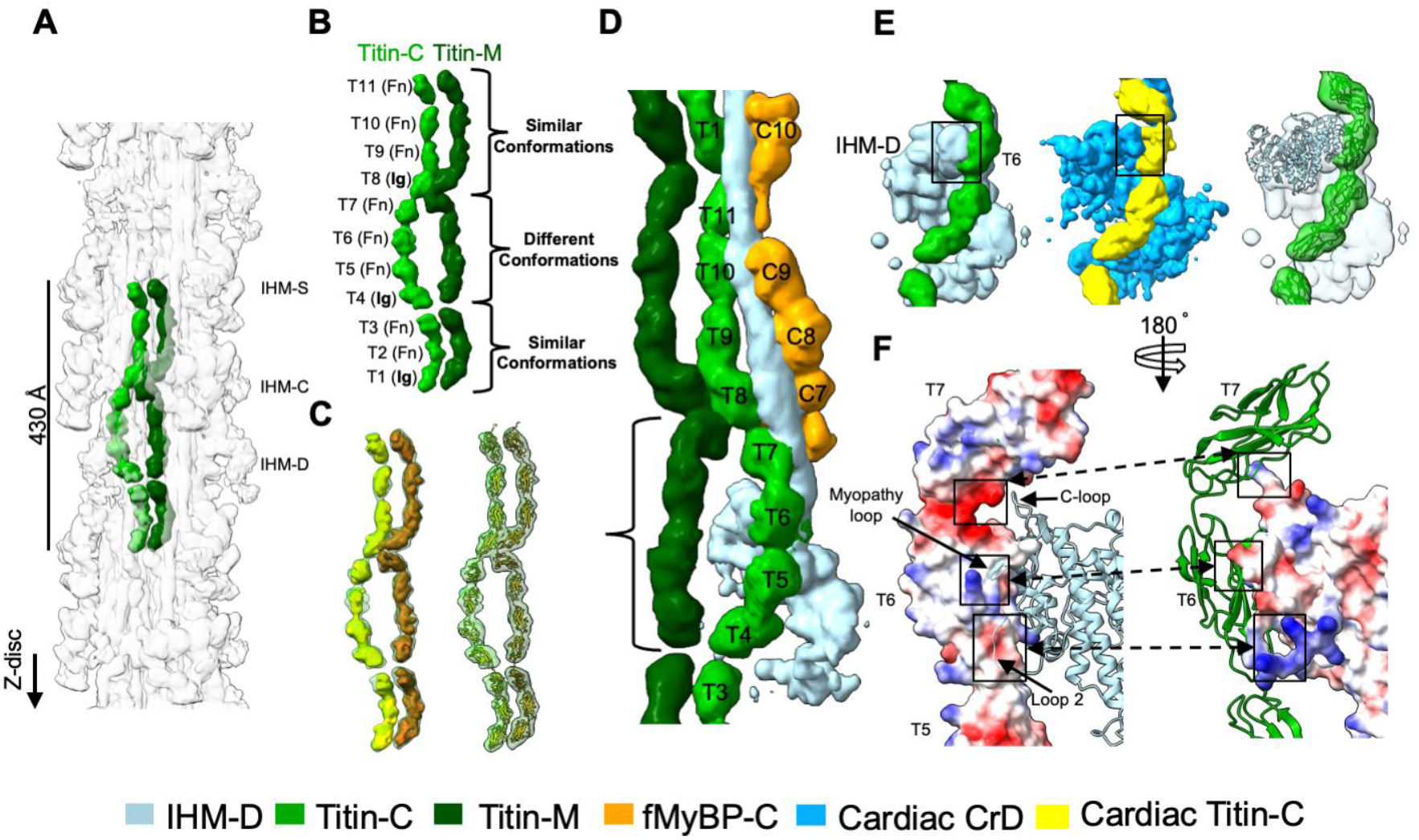
| Titin architecture and domain organization within the C-zone. (A) A pair of eleven-domain titin-C (bright green) and titin-M (dark green) C-zone repeats which align precisely with the arrangement of the three myosin IHMs: IHM-C,-S, and-D. (B) A segmented titin 430Å super-repeat is shown with its domains labeled. While most of the configuration of titin-C and titin-M is similar, their T4-T7 domains differ significantly. (C) To highlight the similarities and differences between cardiac and skeletal titins, we fitted the EM model (left) and atomic structure of cardiac titin (Titin A: ochre, Titin B: yellow; right) from PDB 8G4L into our map. The fit is almost exact resulting in a global change in the shade of green for the two skeletal titins. (D) As in B, the two titin strands adopt distinct conformations; in the T4–T7 region, Titin-C bends away from the Titin-M path toward the myosin backbone to contact the proximal S2 segment and motor domain of IHM-D. Brackets mark the region of different titin conformations. (E) While the IHM-D signal is weak due to its disordered nature, a clear free head myosin density (light blue) is observed at the T6 domain of Titin-C (light green). This interaction pattern is also evident in cardiac titin, as shown in the middle panel. The final panel displays an atomic model of titin-C and IHM-D fitted into the density map of the left panel. (F) The left panel shows titin as an electrostatic surface (blue positive; red negative) with the myosin atomic model. The right panel shows the reciprocal view with the myosin electrostatic surface and titin atomic model. The figure highlights three myosin loops (Loop 2, the C loop, and the myopathic loop) that align with the T6 domain of titin-C. See also Supplementary Video 3.

Similar to the cardiac thick filament^6-8^, we observe distinct regions of conformational similarity between the paired titin-C and titin-M strands (Fig. 3B). The domains at the ends of the super-repeat (T1-T3 and T8-T11) adopt similar conformations in both strands. In contrast, the central segment, comprising domains T4-T7, exhibits significantly different conformations between the two strands. The paired titin conformations observed here are nearly identical to those observed for titin in human cardiac thick filaments^7^ (Fig. 3C).

The organizing role of the titin strands is further clarified by mapping the positions of the myosin heads and fMyBP-C relative to the titin super-repeat (Fig. 3D). Our map shows that fMyBP-C binds exclusively adjacent to domains T8-T11 and T1 of the next titin super-repeat, the region where the Titin-M and Titin-C strands are parallel. Conversely, no MyBP-C density is found along the central T4–T7 region, where the conformations of the titin strands diverge (Fig. 3D).

The direct evidence for this structural conservation is observed when the human cardiac titin structure (from PDB 8G4L)^7^ is fit into our density map of skeletal muscle titin (Fig. 3C). The fit is exceptionally precise. The two cardiac titin strands (Titin A, orange; Titin B, yellow) trace the paths of the skeletal Titin-C and Titin-M densities with high fidelity. This alignment demonstrates that the overall molecular architecture, including the path of the polypeptide chain, the arrangement of the eleven constituent domains, and the specific inter-domain angles and kinks that allow the super-repeat to conform to the 430Å periodicity, is virtually identical between skeletal and cardiac muscle. This conservation of the fundamental “beads-on-a-string” structure, including the kinking pattern that accommodates the 11 domains within the myosin head repeat, indicates that the principles governing titin’s role as a molecular ruler are universal across vertebrate striated muscles^7^.

Although the IHM-D heads are largely disordered in our reconstruction, our map reveals a consistent and clear bridge of density connecting titin T6 to the surface of the IHM-D head region (Fig. 3E). This feature is present at each 430Å repeat and follows the same axial register along the filament. To test whether this density is reproducible in another vertebrate systems, we examined the human cardiac thick filament map (EMD-29722)^7^. Segmentation of titin and the IHM-D heads in that map shows a similar patch of density extending from the titin strand toward the proximal surface of the heads at the position corresponding to titin T6 (Fig. 3E), indicating that this feature is repeatable, is unlikely to arise from noise, and is conserved between skeletal and cardiac thick filaments.

To identify the myosin elements that approach titin T6, we docked the IHM atomic model and the titin C-zone super-repeat into the skeletal filament density and calculated electrostatic surface potentials (Fig. 3F). Because the global resolution of our map is approximately 10Å, we did not attempt de novo modeling of side-chain contacts and instead used rigid-body fitting of existing atomic models to define the interface^32^. Displaying the electrostatic surface on titin with the myosin head shown as an atomic model reveals a negatively charged region on titin T6 opposite a positively charged region on the head that includes loop 2, the C loop, and the myopathy loop of myosin. The complementary electrostatic surfaces at this location identify titin T6 and these three head loops as the most likely contributors to the density that links titin to the IHM-D heads (Fig. 3F).

Titin interactions with myosin tails is complex (Fig. 4A). In the extended 19-crown reconstruction, which resolves the full length of the three tail types, tails from IHM-D run near the filament surface with almost no direct contact with Titin-C and only occasional approaches to Titin-M near the proximal S2 region (Fig. 4B,C). Tails from IHM-S follow a trajectory closer to the filament core and pass near Titin-M but show only very minor contact and essentially no contact with Titin-C (Fig. 4D,E). In contrast, tails from IHM-C remain closely opposed to the titin strands for most of their length, forming an extended series of contacts that are especially prominent with Titin-M (Fig. 4F,G). This hierarchy of interactions, with the strongest and most continuous contacts with titin formed by the IHM-C tails, is similar to the titin and myosin tail network described for the human cardiac thick filament^6-8^.

**Fig 4.**
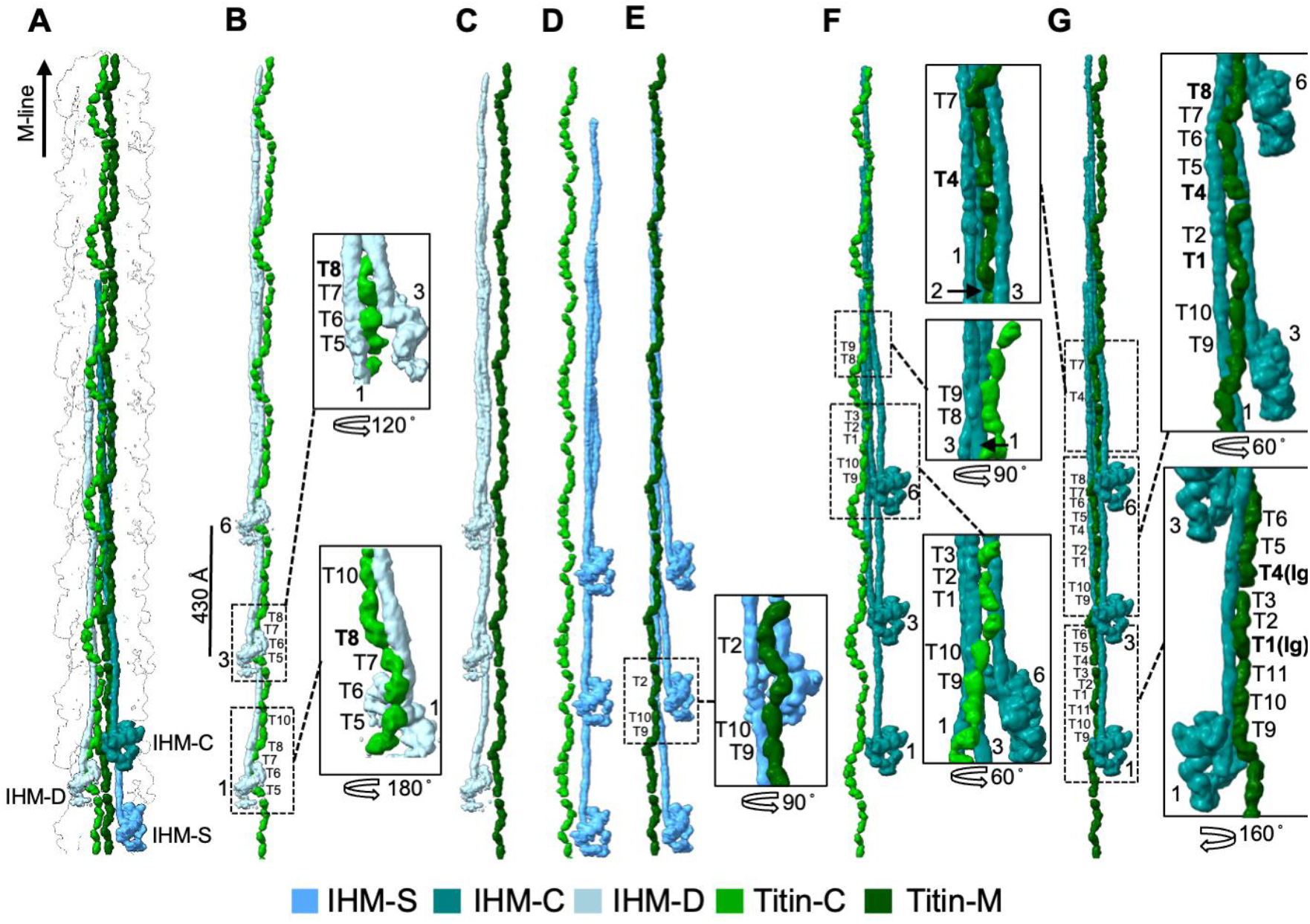
| Titin-Myosin interfaces dictate thick filament organization. (A). Density map showing one pair of titin strands (titin-C: bright green, and titin-M: dark green) and myosin molecules IHM-D, IHM-C, and IHM-S, to show their molecular organization relative to the titins. (B-G) Detailed views of all interactions between individual titin strands and myosin tails. T-numbers indicate the individual titin domains involved (T1-T11) and numbers alone indicate crown portion of myosin tails (crowns 1, 3, and 6, counted distally). (B) IHM-D tail binds to two repeats of the T5-T8 domains of Titin-C (Ig–Fn–Fn–Fn), which are in a distinct conformation relative to Titin-M (Fig. 3B). (C) IHM-D tail does not interact with Titin-M. (D) IHM-S tail does not interact with Titin-C. (E) IHM-S interacts minimally (T2, T9, and T10 domains) with Titin-M, showing a weak but present interaction. (F) Only the LMM segment of IHM-C interacts with Titin-C. (G) IHM-C tail extensively interacts with titin-M. T7 and T8 are the only two domains of Titin-M that do not bind to proximal S2 of the tail of IHM-C.

### fMyBP-C engages IHM-D tail, IHM-S, IHM-C heads, and Titin

We resolved six domains of fMyBP-C, C5-C10. The positions of C7-C10 correspond to the positions found in the cardiac thick filaments^6-8^. Locations of C5 and C6 are different from the cardiac thick filament.

Previous studies suggest that in the sarcomere, while the C-terminal half of fMyBP-C runs parallel to the thick filament, the N-terminal domains of fMyBP-C extend away from the thick filament to bind to the thin filaments^6,8,33^. The six C-terminal domains of fMyBP-C (C5-C10) are visible in the globally averaged map (Fig. 5A); the remaining N-terminal domains are disordered, resulting in no signal because, in the absence of thin filaments the N-terminal half of fMyBP-C has nothing to bind.

**Fig 5.**
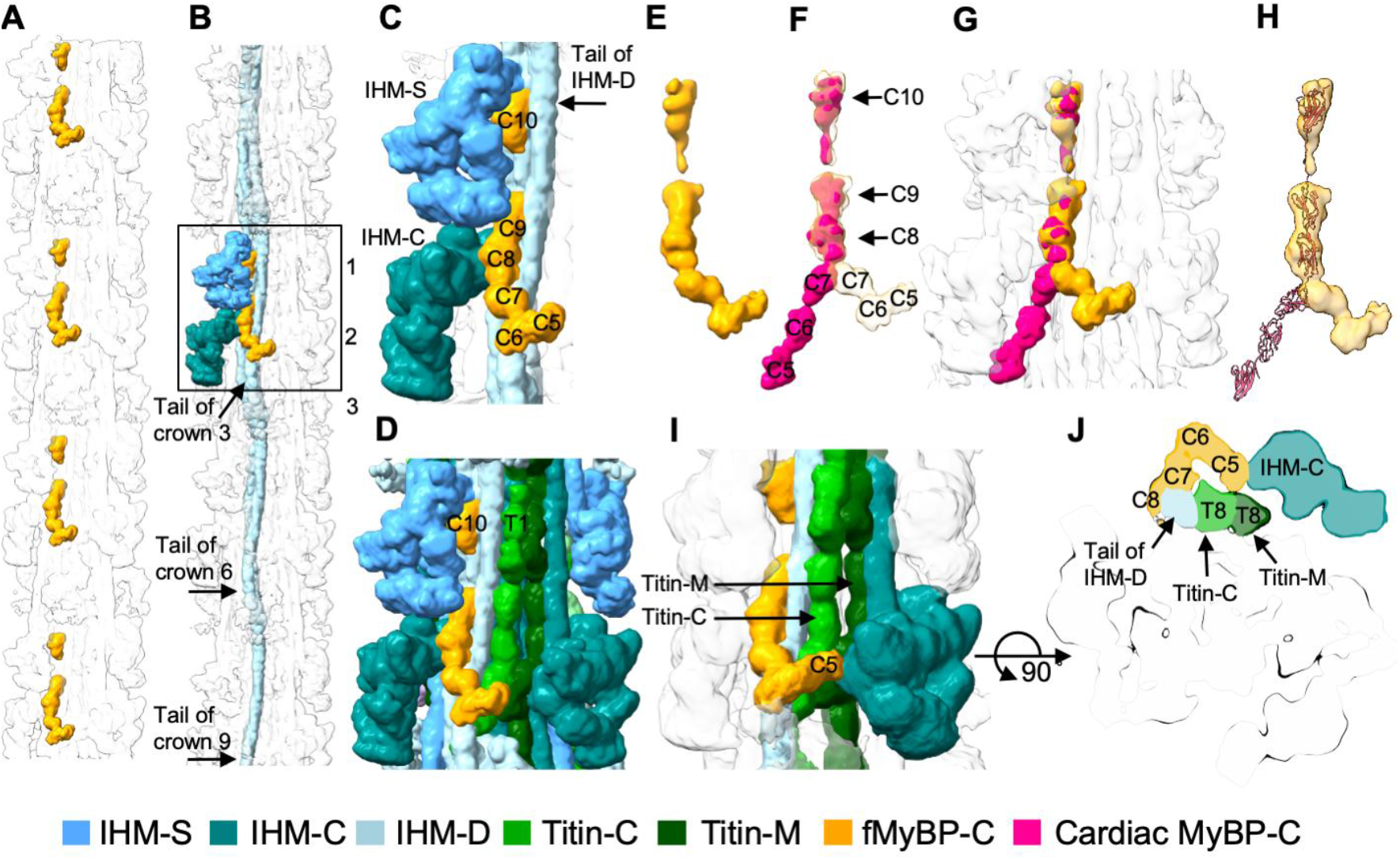
| MyBP-C engages IHM-D tail, IHM-S, and IHM-C heads. (A) 11-crown map provides a comprehensive look at the organization of fMyBP-C (Orange), showing its placement at 430 Å intervals within one segment of the thick filament C-zone. (B) The map further details the interactions between fMyBP-C and myosin molecules (enclosed in a box), specifically highlighting contacts with both the IHM-S and IHM-C heads as well as a sheet of tails derived from three crowns of IHM-D (crowns 3, 6, and 9). (C-D) Magnified views of the map reveal the interactions occurring between the C8-C10 domains and the IHM-D tails. (E) The density of fMyBP-C (colored yellow). (F) The density for domains C10 to C5 of cardiac MyBP-C (colored red) is fitted into our map. (G-H) Highlight similarities and differences between cMyBP-C and fMyBP-C, the map and atomic structure of cardiac MyBP-C (red) are fitted into the fMyBP-C map (orange). All cardiac MyBP-C maps and atomic models referenced above are sourced from PDB 8G4L. (I-J) At a lower contour threshold transverse view and side view a significant density for domain C5 is revealed that is not observed at a higher-contour cut-off; C5 appears to bind to the T8 domain of Titin and the neighboring IHM-C head motif. See Supplementary Video 4 for 3D details.

In our thick filament reconstruction, fMyBP-C domains C8-C10 appear attached to the backbone surface at a location covered by the IHM-D tail layer with C8 contacting the IHM-C free head and the myosin tails (Figs. 5B,C,D). The conformations of C8–C10 are similar in all reports by cryoEM of cardiac MyBP-C structure^6-8^, but differences are observed in the positions of domains C5-C7 (Figs. 5E,F). We compared the conformation of fMyBP-C of this work with its counterpart from the isolated cardiac thick filament by fitting the map (EMD_29726) and atomic models (PDB 8G4L) into the density map (Fig. 5G,H). The comparison reveals that the C8-C10 domains of both structures are similar (Fig. 5F). However, our density for C7 indicates a direction away from the thick filament thereby differing from the single-particle cardiac study while aligning with other published reports on *in situ* cMyBP-C from cryoelectron tomography^6,8^.

Binding studies indicate that titin anchors cMyBP-C to the thick filament^31^. This happens when cMyBP-C’s C-terminus (C8-C10) interacts with the first Ig domain of titin’s 11-domain super-repeat. In our reconstruction we see that C10 of fMyBP-C in the reconstruction aligns with T1 of titin and C8 and C9 extend over T9 and T10 but they are clearly separated by an IHM-D tail (Fig. 4C and D)

At a lower contour cut off than described above, clear but weaker density is seen for possible contact between fMyBP-C domain C5 with the T8 domain of Titin-C/Titin-M and the neighboring IHM-C head motif (Figs. 5I,J). Notably, the change in direction of the C5-C6 domain of fMyBP-C compared with cMyBP-C aligns precisely with T8 of titin. The two titins also interact in this location, and their structure is elevated from the backbone myosin tails and make it possible to interact with non-myosin proteins. These observations show that fMyBP-C and the titins have interactions but not at the C-terminus end of fMyBP-C. Instead, the interaction is at the fMyBP-C C5 domain.

In summary, our findings reveal that while its C-terminal domains (C8-C10) directly attach to the thick filament backbone, its interaction with titin appears to occur via the C5 domain rather than the C-terminus, which is a key difference from cardiac MyBP-C^7^.

## Discussion

The first attempt at imaging isolated vertebrate striated muscle thick filaments was carried out on rabbit psoas muscle thick filaments in 1963^34^, which established that the filaments were bipolar, 1.6 μm in length with a central bare zone of 1600Å and myosin heads projecting from a relatively dense backbone. At that time little was known of the protein composition, other than the filaments contained myosin; MyBP-C and titin had not yet been discovered. A model for the myosin arrangement into “curved molecular crystalline layers” was proposed in 1973^35^. While generally correct for myosin filaments from invertebrates, for which the assumption that myosin was helically arranged proved correct^9^, turned out to be only partially correct for vertebrate striated muscle thick filaments, for which the filaments are not even quasi-helical. Fast forward 60 years to 2023, and cryoEM^7^ and cryoET^6,8^ have provided subnanometer resolution reconstructions of human and mouse cardiac muscle thick filaments, showing in detail the variability in the myosin head and tail arrangement as well as the position, and possible role of the myosin accessory proteins. The differences between the human cardiac thick filament structures can plausibly be explained as differences between isolated and *in situ* thick filaments and the use or not of mavacamten. However, the isolated skeletal thick filament structure shown here was obtained using mavacamten and is thus directly comparable with the isolated cardiac counterpart.

### Shared Architectural Principles: The Conserved Thick Filament Structure Across Skeletal and Cardiac Muscle

A striking feature of the rabbit skeletal muscle thick filament is its high similarity with both human and mouse cardiac muscle, even though these two types of striated muscle function somewhat differently. A skeletal muscle can adjust its amount of force by activating different motor units within a muscle or muscle group (recruitment) and the frequency of motor neuron firing (rate coding)^36,37^. In contrast, cardiac muscle functions as a syncytium where all cardiomyocytes participate in every contraction. Therefore, the strength of contraction must be modulated by mechanisms intrinsic to the cell rather than by recruitment. In cardiac muscle, the force generated varies on a beat-to-beat basis. The volume of blood returning to the heart dictates the degree of stretch and, consequently, the force generated by the cardiomyocytes (Frank-Starling law). While cardiac muscle relies heavily on these beat-to-beat changes in sarcomere length to modulate force, skeletal muscle operates within a length range defined by joint mechanics. Hence, subtle structural differences in the thick filament may underlie the distinct activation mechanisms found in cardiac versus skeletal muscle. On that basis it is perhaps not surprising that the thick filaments of the different muscle types are so similar even among widely different animal species.

Cardiac muscle’s distinctly different regulatory problem is solved in part by a less cooperative response on the part of its thin filament to calcium concentration in which the amount of calcium released produces different degrees of the thin filament activation^38^. The skeletal muscle thin filament has higher cooperativity and thus a more complete thin filament activation in combination with higher Ca^2+^ concentration achieved during a shorter duration Ca^2+^ transient^39,40^. Efficiency would suggest that the degree of cardiac thick filament activation should be tuned to the degree of thin filament activation^15^, a role that in part may be played by MyBP-C.

### Myosin-Binding Protein C: Distinct Interactions with Titin and Myosin Tails

The greatest, though still small, difference between the isolated cardiac and skeletal thick filaments is the position of the middle domains of fMyBP-C. In the isolated cardiac thick filaments, lacking the presence of thin filaments, the middle domains of cMyBP-C were tucked into a space that contacted the myosin regulatory light chain and proximal S2^7^. Here fMyBP-C takes an alternative route and instead seems to contact titin and IHM-C. In both cases, lacking the appropriately placed thin filament enhanced this distinctive alternative binding site.

Evidence for a titin-MyBP-C interaction comes from early biochemical and structural experiments in which C-terminal fragments of MyBP-C were shown to bind specific Ig domains within titin C-zone super-repeats^31^, and from mapping of these titin domains to the C-region of the thick filament^41^. Immuno-electron microscopy and more recent cryoEM studies further place the C-terminal MyBP-C collar along titin’s C-zone super-repeats on the surface of the thick filament^42,43^. Our skeletal thick-filament structure extends this picture by revealing that the only direct contact we can detect between MyBP-C and titin occurs at a more internal position, where C5 approaches titin domain T8, a location in agreement with earlier studies^31^. In agreement with other cryoEM structures, the C-terminal domains C8–C10 lie close to titin but are clearly separated from it by the IHM-D tails. Thus, in fast skeletal muscle, titin appears to anchor the central portion of MyBP-C rather than its extreme C-terminus, creating a single, well-defined linkage between the titin filament and the MyBP-C collar in each C-zone repeat.

We would note that domain C5 of fMyBP-C has a much shorter CD loop, lacking the cardiac-specific 28-amino-acid insert^44^, and this structural difference is likely central to the distinct titin-binding behavior we observe. NMR and computational studies have shown that this extended CD loop in cardiac C5 is highly dynamic, largely unstructured and destabilizes the isolated C5 domain, in contrast to other, more stable Ig-like domains of titin and MyBP-C^44,45^. Doh et al. directly tested this idea by expressing C4-C5 constructs with and without the linker and loop and showed that the linker and C5 loop modulate both secondary structure and thermal stability of the tandem: deleting either region, especially the linker, shifts the melting behavior toward higher temperatures and changes the accessible conformations. Their molecular-dynamics “hinge-and-latch” model proposes that the 10-residue linker provides flexibility, while the cardiac-specific C5 loop acts as a latch that helps C5 fold back onto C4 and stabilizes a sharply bent V-shaped conformation enriched in cardiac, but not skeletal, MyBP-C. In rotary-shadowing experiments, skeletal MyBP-C molecules, lacking this loop, indeed populate a more open U-shaped conformations with larger hinge angles, consistent with a less tightly latched C4-C5 segment.

In our fast skeletal thick filament, which lacks the destabilizing C5 loop, the C4-C5 region is expected to resemble the loop-deleted state, with a more stable C5 domain and a hinge that is not forced into the tightly bent, latched configuration^45^. This provides a natural explanation for why C5 in fMyBP-C is able to pivot away from the myosin neck region and toward titin T8, forming the single titin contact that we observe in the super-relaxed–like configuration, whereas no equivalent C5–titin contact has been resolved in cardiac thick-filament structures. In cardiac muscle, the long, unstable C5 loop may bias C4-C5 toward a compact, V-shaped conformation that favors interactions near the regulatory light chain and effectively prevents C5 from reaching a T8-equivalent titin site. A C5–T8 interaction in fast skeletal muscle would mechanically couple the compliance of titin’s C-zone super-repeat directly to the fMyBP-C collar and, through fMyBP-C, to the folded myosin heads. This arrangement provides a structural route by which titin strain in this region could contribute to thick filament mechanosensing^15^ and decrease the fraction of heads in the interacting heads motif and thereby tune contraction speed and energetic cost.

### The Action of Mavacamten Beyond Its Cardiac Indications

Mavacamten is a compound selected for its specificity of inhibiting cardiac myosin^25,46^. Inhibition is believed to occur by promoting myosin heads conforming to the interacting heads motif^26^. Mavacamten is now used in the treatment of obstructive hypertrophic cardiomyopathy^47^. Previous cardiac thick filament structures used mavacamten at a concentration of 20-50 μM^6,7^ which is ∼30x to 80x its reported IC_50_ for human cardiac myosin (IC_50_ 0.58 ± 0.07 µM)^48^. Mavacamten has similar biochemical effects on skeletal muscle^49^ but in our study required ten times as much as the reported K_d_ (14.4±3.2 μM) to rabbit psoas myosin^50^. The binding affinity of mavacamten could be affected by only small changes in its binding site and this is presumably the effect that we observe^27^. Thus, we were able to use mavacamten in our fast skeletal muscle preparations.

### Evolution of the Striated Muscle Thick Filament

The striated muscle myosin filaments of invertebrates are highly heterogeneous, having highly variable lengths^51^, rotational symmetries ranging from 4^9,52^ to 7^53^, and variable amounts of the protein paramyosin^51,54^. Among the specialized thick filaments of insect indirect flight muscle, the filaments vary in length^50^, paramyosin content^54^, ordering of the myosin heads, conformation of the protein flightin^13^, and the presence (or absence) of other non-myosin proteins, e.g. stretchin-klp^13^. The one feature held in common among these filament structures is the arrangement of myosin tails into “curved molecular crystalline layers”^33^. Because of their highly regular helical structure, we have detailed knowledge of the molecular arrangements within indirect flight muscle thick filaments.

Recent phylogenomics has suggested the four insect orders that developed indirect flight muscle, e.g. Hemiptera, Coleoptera, Hymenoptera and Diptera, evolved over 290 – 160 Million years ago (Mya) from a common ancestor that existed 374 Mya^55^. Though evolving from a common ancestor, each insect species whose thick filament has so far been studied evolved a different filament structure presumably to obtain properties appropriate to indirect flight muscle which must contract at a narrow range of high frequencies governed by the deformation mechanics of the thorax. Indirect flight muscle must also support the thoracic endothermy needed to sustain the high ATPase rates needed for high frequency wing beating^16^.

Vertebrates appeared as long ago as 525 – 520 Mya^56^ and from these beginnings, the largest animals to have lived on this planet have evolved. Invertebrates, having appeared probably not that much earlier have also radiated widely in this time period but never evolved into creatures as large as the vertebrates. While invertebrate evolution may have been limited by their exoskeleton among others, it may also be limited by evolution of their striated muscle.

All placental mammals are reported to have evolved from a common ancestor, a shrew-like creature, that lived 102 ± 12 Mya^57^ during the late Cretaceous period (145.5 Mya - 66 Mya). After the K-Pg extinction that eliminated the dinosaurs, these mammals were freed to evolve into the wide range of species, many of which still exist. Among these are the largest species that ever existed on this planet, especially when counting the extinct Dinosauria clade. Rabbits, mice and humans diverged from a common ancestor that lived ∼80 Mya, before the K-Pg extinction^57^. Mice that have been studied extensively first appeared about 5 Mya^58^. These species differ widely in size, though proportionally to about the same extent as the insects with indirect flight muscle that have been studied.

Mice and humans share the same common ancestor while differing greatly in size, and yet their cardiac thick filament structures proved nearly identical within the C-zone, the most homogeneous segment of the vertebrate thick filament. The skeletal muscles of rabbits belong to the other vertebrate striated muscle type. With the possible exception of a single difference in the MyBP-C placement of domain C-5, the C-zone structure of rabbit skeletal muscle thick filaments is almost identical to those from mouse and human cardiac muscle. Fishes evolved very much earlier than placental mammals, yet their muscles appear to be highly similar to those from mammals except for a more ordered arrangement of the thick filaments^59^. Fish, however, are not predominately endothermic.

The thick filament structure of vertebrate striated muscle appears to be highly adaptable to the varying requirements of evolving vertebrate species and different muscle types, i.e. cardiac and skeletal. Conversely, the indirect flight muscles of insects need only contract at a single, or narrowly defined frequency. Yet this variation seems to have required evolution of a different thick filament architecture each time it occurred. For birds and mammals, skeletal muscle plays a central role in endothermy by coupling high rates of ATP turnover and Ca^2+^ cycling to heat production, whereas indirect flight muscle contraction is optimized for mechanical efficiency and power output, with only localized and transient thermogenesis. The vertebrate thick filament’s utility appears to be its adaptability for the variable recruitment of myosin heads for force production during contraction^15^ and the sequestration or release of myosin heads (the interacting heads motif) for energy conservation when relaxed or when producing endothermy^60^.

## Methods

### Sample preparation

All protocols and experimental procedures using animal subjects followed NIH guidelines and were approved by the Florida State University Institutional Animal Care and Use Committee (IACUC), protocol number TR202300000011. Glycerinated rabbit psoas muscle, stored in a low calcium solution (pCa 8) at −20 °C in 50% glycerol, was used as the source of native thick filaments. For each preparation, a small piece (∼2 mm^3^) was cut with a razor blade and transferred to 3 ml relaxing solution (100 mM KCl, 20 mM MOPS, 5 mM EGTA, 5 mM dithiothreitol, 5 mM MgCl_2_ and 5 mM ATP, pH 7.0) at 4 °C to wash out the glycerol. Muscle strips were then gently teased into very thin bundles (0.5–1 mm) and incubated for 30 min at room temperature in relaxing solution containing 0.5% Triton X-100 on a rocker.

Elastase treatment was used to release thick filaments^61^. Elastase (100 µl) was added to 2 ml of sample for 3 min at room temperature, after which 20 µl of 0.1 M PMSF (1 mM final) was added and the preparation was incubated for a further 10 min to stop the reaction. The sample was gently sheared with a 20 G needle to release thick filaments while minimizing breakage. Large tissue fragments were removed by centrifugation at 3,000 × g for 5 min. The supernatant (2 ml) was mixed with recombinant calcium-insensitive gelsolin (2 mg/ml) and incubated for 1 h at room temperature to remove sarcomeric actin^62^. The gelsolin construct was expressed in *Escherichia coli* and purified by standard techniques^63^. Thick filaments were pelleted by centrifugation (14,000 rpm, 5 min) and gently resuspended in 200 µl fresh relaxing solution. Quality and concentration of the thick filaments were assessed by negative-stain EM with 2% uranyl acetate^64^. For mavacamten treatment, 60 µl of the final thick-filament suspension was transferred to a small tube and 0.91 µl of 10 mM mavacamten stock was added to give a final mavacamten concentration of 150 µM.

UltrAuFoil R1.2/1.3, 300-mesh gold grids were plasma-cleaned in an O_2_ mixture for 20 s and with added specimen plunge-frozen on a Vitrobot Mark IV into liquid ethane. Frozen grids were stored in liquid nitrogen and pre-screened on a Titan Krios G1 (FSU Biological Science Imaging Resource) equipped with a DE-Apollo direct electron detector to assess ice thickness, filament integrity, and filament distribution.

### Data acquisition

Movies were collected at NCCAT on a 300 kV Titan Krios equipped with a Thermo Fisher Scientific Falcon IV direct electron detector and a Selectris energy filter. Data were recorded at a nominal magnification of 130,000×, corresponding to a calibrated pixel size of 0.927Å/pixel. A defocus range of −1.0 to −3.0 µm was used, with a total exposure of 48.87 e^−^/Å^2^ per movie. Data were acquired in three sessions and combined for processing, yielding 56,189 movies in total, of which only 17,280 containing well-dispersed, traceable filaments were retained for downstream processing, as most movies either lacked visible thick filaments or exhibited extensive filament overlap. Facility-standard motion correction, Motioncor2,^65^ was performed at NCCAT prior downloading data.

### Data processing

All processing was performed in cryoSPARC v4.6.2 (Structura Biotechnology)^66^. Patch CTF was used for per-micrograph CTF estimation. For particle identification, a small set of filaments was first manually picked to generate high-quality 2D class templates, which were then used to guide automated picking with the cryoSPARC Filament Tracer. Segments were extracted in 1120-pixel boxes and Fourier-cropped to 400 px (effective pixel size ≈ 2.60Å/pixel) for classification. Multiple rounds of reference-free 2D classification removed contaminants and empty boxes, yielding 134,840 high-quality segments. An ab-initio 3D reconstruction was generated and followed by heterogeneous refinement into four classes; two classes were retained for further analysis (35,406 and 43,696 segments, respectively), while the remaining two were discarded. Each retained class underwent non-uniform refinement (no additional beam-tilt/optics-group refinements applied)^67^, producing maps at ∼10Å global resolution (FSC 0.143). For visualization, maps were post-processed with DeepEMhancer^68^, and local-resolution estimates are provided in the Supplementary Information. Figures were made using UCSF ChimeraX v1.11^69^.

## Acknowledgements

Supported by NIH grants R21 AR077802 (to J.R.P. and K.A.T.), R35 GM139616 (to K.A.T.) and R01 HL160966 (to J.R.P.). J.R.P. acknowledges support from the Institute of Pediatric Rare Disease at Florida State University College of Medicine. Some of this work was performed at the National Center for CryoEM Access and Training (NCCAT) and the Simons Electron Microscopy Center located at the New York Structural Biology Center, supported by NIH (Common Fund U24 GM129539, R24 GM154192), the Simons Foundation (SF349247), and the NY State Assembly. Some microscopy was carried out at the Biological Science Imaging Resource of Florida State University supported partially by NIH grant R24 GM145964. We thank Belinda Bullard and P. Bryant Chase for their comments on an earlier draft.

