## Supplementary text for "Myosin Filaments of Vertebrate Skeletal and Cardiac Muscle are Highly Similar, but not Identical"

### Supplementary Figure

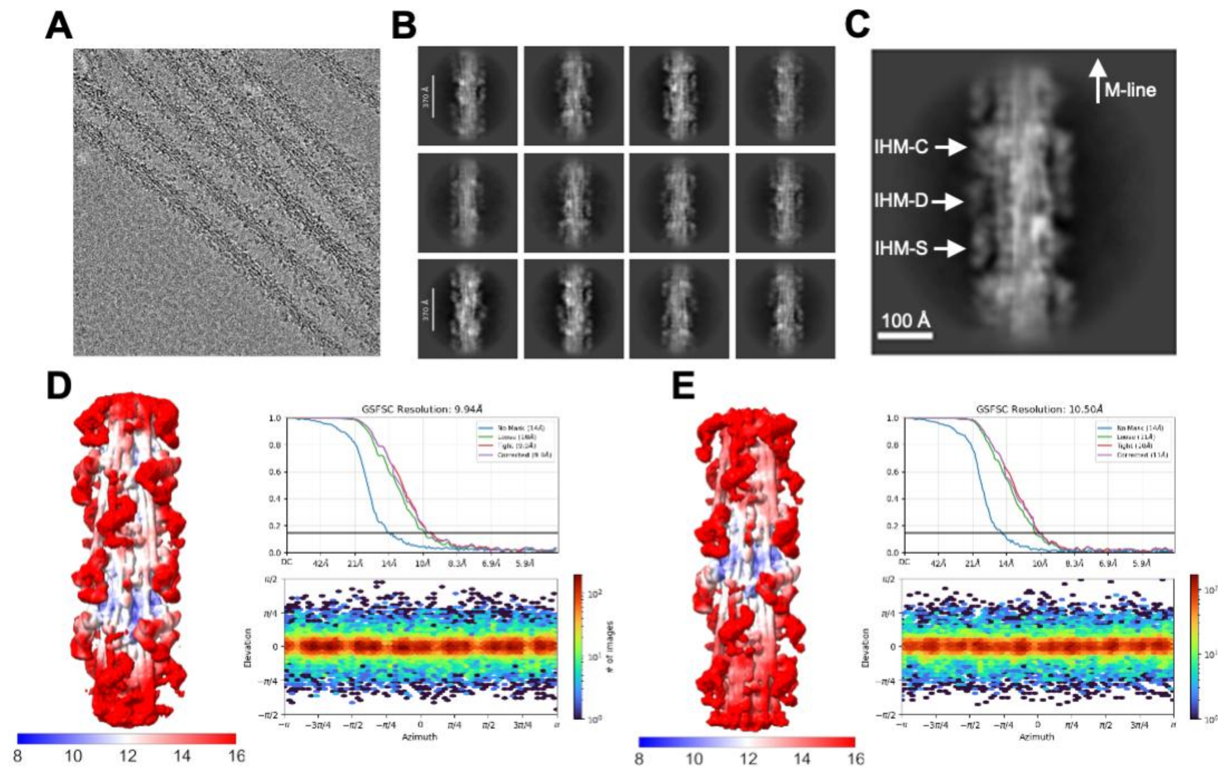


**Extended Data Fig. 1 |The process of cryo-electron microscopy (cryo-EM) imaging and the subsequent data processing.**

(A) cryo-EM image of a thick filament from the rabbit psoas muscle. (B) This was followed by the generation of two-dimensional (2D) class averages from segments of the C-zone, each seven crowns in length. (C) An enlarged view of a 2D class average reveals that two of the three interacting heads of the myosin molecule (IHM-S and IHM-C) exhibit strong density, whereas the third (IHM-D) appears diffuse. This suggests that the third head is mobile. (D) The two 7-crown reconstructions are presented with colored maps that depict the local resolution in Angstroms, as indicated by the scale bar. The global resolution estimate for each reconstruction is provided by gold-standard Fourier shell correlation (GSFSC) curves, positioned in the upper portion of the figure. Additionally, the heat maps situated at the bottom illustrate the distribution of viewing angles of the particles that were

### Supplementary Videos

#### Supplementary Video 1

Overall architecture and segmentation of the 7-crown thick-filament map. The movie opens with the 7-crown cryo-EM reconstruction of the rabbit psoas thick filament, sharpened using DeepEMhancer, shown rotating continuously about the y-axis to provide a global three-dimensional view. After a 180° rotation, the segmented map is introduced to delineate the major structural components of the filament. Following a further 180° rotation, the view zooms in on a three-crown region corresponding to a single axial repeat, revealing finer structural detail and local organization within the filament. Color scheme: IHM-D (light blue), IHM-S (blue), IHM-C (teal), Titin-M (dark green), Titin-C (bright green), and fMyBP-C (orange).

#### Supplementary Video 2

19-crown map showing distinct myosin tail layers. The 7-crown reconstruction is extended to a 19-crown segmented map to enable tracking of myosin molecules across multiple axial repeats. The extended map is rotated about the y-axis, revealing the organization of full-length myosin tails within the thick-filament backbone. The left panel shows the three myosin classes together (IHM-D, IHM-S and IHM-C), illustrating the combined organization of one repeating unit. The remaining panels show the individual tail layers corresponding to IHM-D, IHM-C and IHM-S, highlighting their distinct radial positions and trajectories within the filament. Color scheme: IHM-D (light blue), IHM-S (blue), IHM-C (teal).

#### Supplementary Video 3

Fitting of a cardiac IHM head into the IHM-D density and visualization of titin–myosin interactions. An interacting-heads motif (EMD-15354 is fitted into the IHM-D density of our map. The reconstruction is shown rotating about the y-axis, with the density displayed semi-transparently to illustrate the placement of the head within the map. The titin-C segment is then introduced, followed by the display of atomic models for titin-C (based on PDB:8G4L) and the myosin motor domain (PDB:2MYS). The visualization concludes with the isolated atomic models, highlighting the interaction between titin T6 and surface loops of the myosin motor domain. Color scheme: IHM (light blue); titin-C (bright green); myosin motor domain (light blue); C loop (blue); myopathy loop (yellow); loop 2 (red).

**Supplementary Video 4**

Low-threshold map highlighting titin–fMyBP-C interactions. A low-contour-threshold cryo-EM map is shown to reveal interactions between titin and fast myosin-binding protein C (fMyBP-C) in the rabbit psoas thick filament. The reconstruction is displayed semi-transparently, with segmented densities for titin, fMyBP-C and the interacting-heads motif overlaid. The map is rotated about the y-axis to provide an overall view, followed by a zoom-in to emphasize the interaction region, and a subsequent rotation about the x-axis to further illustrate the three-dimensional organization of these contacts. Color scheme: IHM-D (light blue), IHM-C (teal), Titin-C (bright green), and fMyBP-C (orange).
